# Contribution of the endosomal-lysosomal and proteasomal systems in Amyloid-β Precursor Protein derived fragments processing

**DOI:** 10.1101/300921

**Authors:** Caroline Evrard, Pascal Kienlen-Campard, Rémi Opsomer, Bernadette Tasiaux, Jean-Noël Octave, Luc Buée, Nicolas Sergeant, Valérie Vingtdeux

## Abstract

Aβ peptides, the major components of amyloid deposits of Alzheimer’s disease, are released following sequential cleavages by secretases of its precursor named the amyloid precursor protein (APP). In addition to secretases, degradation pathways, in particular the endosomal/lysosomal and proteasomal systems have also been reported to contribute to APP processing. However, the respective role of each of these pathways towards APP metabolism remains to be established. To address this, we used HEK 293 cells and primary neurons expressing full-length APP^WT^ or the β-secretase-derived C99 fragments (β-CTFs) in which degradation pathways were selectively blocked using pharmacological drugs. APP metabolites, including carboxy-terminal fragments (CTFs), soluble APP (sAPP) and Aβ peptides were studied. In this report, we show that APP-CTFs produced from endogenous or overexpressed full-length APP are mainly processed by γ-secretase and the endosomal/lysosomal pathway, while in sharp contrast, overexpressed C99 alone is mainly degraded by the proteasome and to a lesser extent by γ-secretase.

## Introduction

Amyloid deposits, that mainly consist of the extracellular aggregation of amyloid β-peptides (Aβ) into plaques, are one of the major pathological hallmarks of Alzheimer’s disease (AD) and the main targets of current clinical trials [1]. Aβ peptides are generated by sequential and compartmentalized cleavages of the type I transmembrane protein named Amyloid Precursor Protein (APP). Proteolytic cleavage of APP by β-secretase (mainly BACE1), characterized as the first step of the amyloidogenic pathway, releases APP carboxy-terminal fragments (APP-CTFs) named β-CTF or C99 that remain membrane bound and the soluble sAPPβ fragments that are secreted. β-CTFs are then processed within its transmembrane domain by γ-secretase, a transmembrane proteolytic complex, thereby releasing Aβ peptides and APP Intracellular Domain (AICD) (for a review see [2]). Alternatively to this amyloidogenic pathway, APP can be processed by an α-secretase activity [3], which cleaves within the Aβ sequence thus precluding its production. This α-secretase cleavage of APP releases soluble sAPPα fragments that are secreted and membrane-bound APP-CTFs named α-CTFs or C83, that can be further processed by γ-secretase (Fig 1A). Recently, the identification of new secretases, like δ-secretase (asparagine endopeptidase), η-secretase (MT5-MMP) and meprin β, have added to the complexity of APP processing [4]. Meprin β can be considered as an alternate β-secretase enzyme since it gives rise to a β-CTF starting at position 2 [5,6]. δ-secretase and η-secretase generate larger APP-CTFs that can be further processed by α- or β- and γ-secretases [7–9] (Fig 1B). Finally, it is also important to note that these secretases are active at various intracellular locations. While their exact location is still debated, we propose a schematic representation of their spatial location for APP cleavage in agreement with literature (Fig 1C).

**Figure 1:**
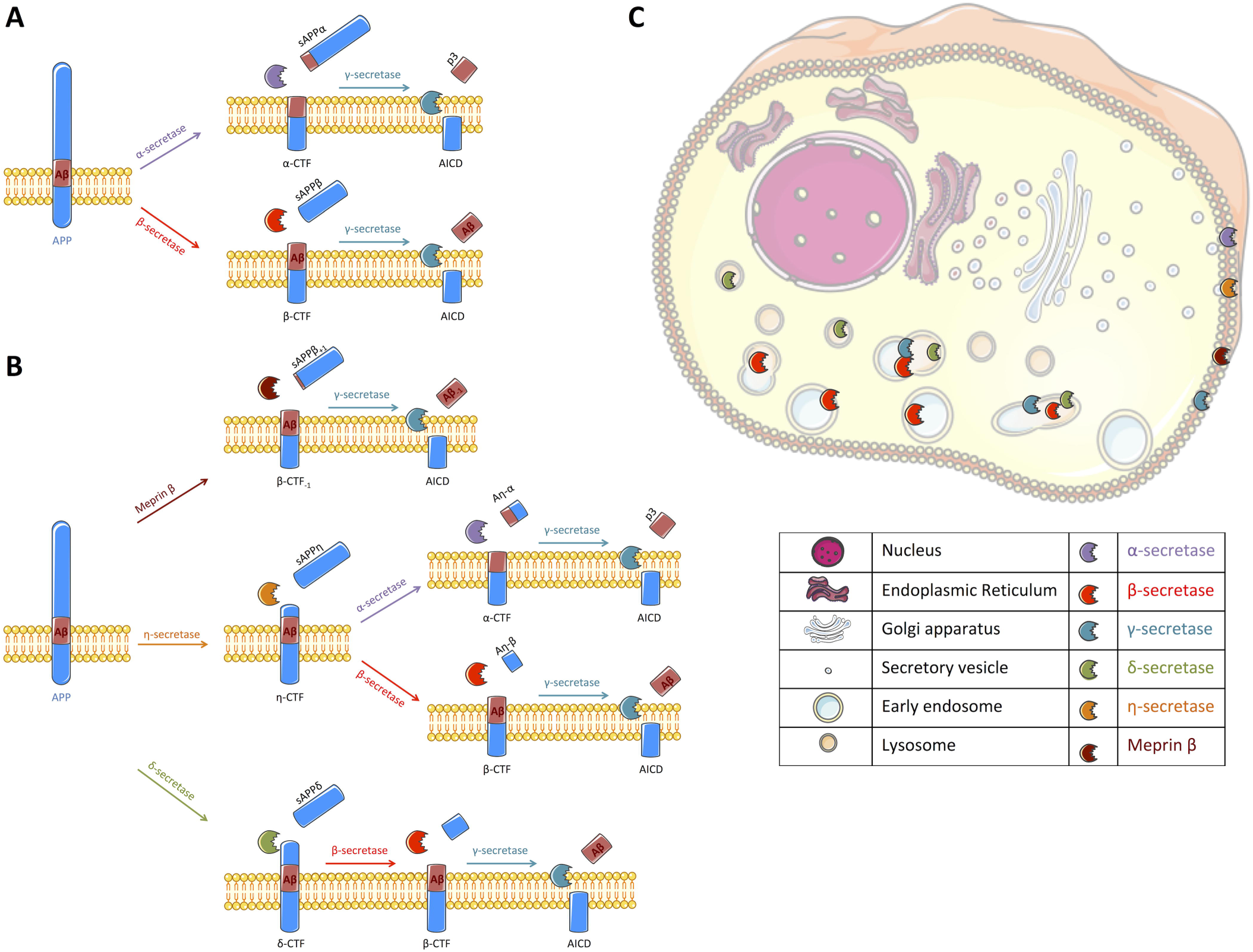
Schematic representation of APP processing and the cellular localization of the different secretases involved. **(A)** Following synthesis and maturation, the amyloid precursor protein (APP) can classically follow two different pathways: the non-amyloidogenic pathway or the amyloidogenic, the latest being at the origin of Aβ peptides production. In the non-amyloidogenic pathway, APP is first cleaved by α-secretase (*mallow*) within the Aβ sequence, liberating a soluble fragment sAPPα and a transmembrane carboxy-terminal fragment (APβ-CTF) called α-CTF. Then, γ-secretase (*turquoise*) processes α-CTF into p3 and APP intracellular domain (AICD). In the amyloidogenic pathway, sAPPβ and β-CTF are released following APP cleavage by β-secretase (*red*). β-CTF is further processed by γ-secretase into Aβ peptide and AICD. **(B)** Alternatively to a direct cleavage by α- or β-secretase, APP can be processed by other secretases that have been recently described. Meprin β (*brown*) can act as an alternative β-secretase releasing sAPPβ with 1 amino acid longer and β-CTF with an amino acid less. β-CTF. _1_ is similarly cleaved by γ-secretase. η-secretase (*orange*) cleaves APP into sAPPη and η-CTF. η-CTF can follow either the amyloidogenic or the non-amyloidogenic pathway, releasing Aη- β and β-CTF or Aη-α and α-CTF, respectively. The remaining CTFs (α- and β-) are likely processed by γ-secretase. δ-secretase (*green*) processes APP into sAPPδ and δ-CTF. Cleavage of δ-CTF by ̡- and γ-secretase releases Aβ and AICD. **(C)** All these secretases are active in different cellular compartments that remain debated in literature. Thus, we propose a schematic representation of their spatial location for APP cleavage in agreement with literature, α-secretase and meprin β were described to cleave APP at the cell surface where α-CTF and β-CTF-_1_can then be processed by γ-secretase [2,40,41], γ-secretase cleavage was reported to occur at the plasma membrane and in endosomes [42], β-secretase processing of APP is reported to occur in endosomal/lysosomal acidic compartments as it is the case for the δ-secretase [7,31] where APP-CTFs could also be cleaved by γ-secretase. The location of APP processing by η-secretase is still undefined but it is likely that η-secretase is located at the plasma membrane [43]. Figure was produced in part using Servier Medical Art.

Among all the fragments generated along APP processing, β-CTFs are of particular interest since they are the direct precursors of Aβ peptides. In addition, β-CTFs have been shown to accumulate before Aβ peptides *in vivo*, and suggested to be instrumental in the initiation of the neurodegenerative process and cognitive alterations [10]. Inhibition of β-secretase cleavage of APP rescues long-term potentiation (LTP) and memory deficits in a mouse model of AD [11]. Of note, β-CTFs accumulation was also described to occur in AD brains [12]. While the exact impact of β-CTFs accumulation in AD is not completely elucidated yet, a better understanding of the mechanisms involved in APP-CTFs, and in particular β-CTFs, degradation is needed.

Indeed, alternatively to their cleavage by γ-secretase, lysosomal proteases and/or the proteasome are involved in APP-CTFs degradation. APP has an internalization carboxy-terminal NPxY motif and is thus found along endosomal/lysosomal compartments [13–15]. Inhibition of lysosomal proteases induces an accumulation of APP-CTFs [16–18], showing that the endosome/lysosome pathway is important for the processing of APP. Other reports suggested that APP-CTFs, in particular β-CTFs, were rather processed by the proteasome [19–21]. Additionally, the proteasome was also suggested to indirectly modulate APP metabolism by regulating the degradation of proteases implicated in APP metabolism [22,23].

Although the γ-secretase, the proteasome and the lysosomal proteases were shown to be involved in APP processing and β-CTFs degradation, their mutual implication remains to be established. In the present study, we compared the contribution of these three degradation pathways on APP processing in HEK cells and in rat primary neuronal cultures expressing either APP^WT^ or the C99 [24], by using selective pharmacological drugs.

## Materials and Methods

### HEK cell culture and primary neuronal cell culture

HEK293 stably expressing or not APP^WT^ or C99 were previously described [25] and kindly provided by Pr. F. Checler. Briefly, HEK cells were grown in Dulbecco’s Modified Eagle Medium (DMEM, high glucose, pyruvate – GIBCO by Life Technologies) supplemented with 10% foetal bovine serum, 2 mM L-glutamine, 1 mM non-essential amino-acids and penicillin/streptomycin (GIBCO by Life Technologies) in a 5% CO_2_ humidified incubator at 37°C. Primary cultures of rat embryonic cortical neurons were prepared as previously described. After 6 days of culture, neurons were infected with recombinant adenovirus expressing either APP^WT^ (corresponding to wild-type APP695 isoform) or C99 as described [26]. All animal procedures used in the study were carried out in accordance with institutional and European guidelines and experimental protocols were approved by the Animal Ethics Committee from the Université catholique de Louvain (UCL, Brussels, Belgium).

### Drug treatments

Bafilomycin A1 (Baf_A1_), Compound E (CompE) and DAPT were purchased from Merck Millipore. Chloroquine (CQ) and Lactacystin (Lacta) were purchased from Sigma-Aldrich. Cells were plated into 6-well plates at a density of 500 000 cells per well, 18 hours (h) before drug exposure. Cultures were briefly washed once with warm phosphate-buffered saline (PBS) and then exposed for 6 or 24h to drug-treatments at the indicated concentrations. For primary cultures, treatments were performed at day 4 post-infection. At the end of treatments, the conditioned medium was collected and kept at -80°C until use for Aβ_1-40_, Aβ_1-42_ peptides and sAPPα/sAPPβ dosage by ELISA and electro-chemiluminescence immunoassay respectively. Then, cells were rinsed once with PBS and collected in 100μL of Laemmli buffer (10mM Tris, 20% glycerol and 2% Sodium dodecyl sulfate) using a cell-scraper. The lysate was sonicated for 5 min before measurement of total protein concentration using the Pierce BCA Protein Assay Kit (Thermo scientific) according to the manufacturer’s protocol. Samples were conserved at -80°C before analysis.

### Western blotting

Equal quantity of total proteins (20 μg/lane) was loaded on a 16.5% Tris-tricine or an 8-16% tris-glycine polyacrylamide gel. Tris-tricine SDS-polyacrylamide gel electrophoresis and Western-blotting were performed as described [27]. APP and APP-CTFs were detected with a rabbit APP-Cter-C17 antibody (1/5000) that is raised against the last 17 amino acids of the human APP sequence [27,28]. The β-tubulin antibody (1/1000) and all secondary antibodies coupled with horseradish peroxidase were purchased from Sigma-Aldrich. Results were normalized prior to APP-CTFs quantification by using the β-tubulin signal as a loading control. Quantifications of protein expression levels were performed with ImageJ Software (NIH).

### Aβ peptides quantification

The collected medium was spun at 200g to eliminate cell debris. Secreted Aβ_1-40_and Aβ_1-42_ peptide concentrations in pg/mL were determined using amyloid beta 40 and 42 Human ELISA kits (Invitrogen) according to the manufacturer’s instructions.

### sAPPα/sAPPβ quantification

Conditioned media of collected at the end of experiments was briefly centrifuged as describe above to eliminated cells debris. sAPPα and sAPPβ concentrations were determined using the sAPPα/sAPPβ multiplex kit (Meso Scale Diagnostics, MSD^®^) according to the manufacturer’s instructions.

### Statistical Analysis

Statistical analyses were performed with GraphPad Prism 6 program (GraphPad Software) by using the unpaired Student’s test for pairwise comparisons. Statistical significance was set at ^∗^p<0.05, ^∗∗p<^0.001, ^∗∗∗^p<0.001 and ^∗∗∗∗^p<0,0001. All data are reported as mean ± standard error of the mean (SEM) of at least n=3 experiments.

## Results

### APP-derived carboxy-terminal fragments are mainly processed byγ-secretase and lysosomal proteases

The contribution of pathways responsible for APP-CTFs degradation was investigated using a pharmacological approach. First, naive HEK (HEK^ctrl^) and HEK stably expressing APP^WT^ (HEK APP^wt^) were treated for 6h with the γ-secretase inhibitor Compound E (CompE) [29] and APP-CTFs were analysed by Western blotting (Fig. 2A). In HEK^ctrl^ cells, endogenous APP-CTFs were barely detected in control conditions whereas they were increased after γ-secretase inhibition with α-CTF being the major species observed. A similar profile is observed in HEK APP^WT^, indicating that overexpressed APP is efficiently processed. The major CTFs species observed are α-CTFs as in HEK^ctrl^ cells; showing that there is no bias in the amyloidogenic/non-amyloidogenic balance induced by overexpression. γ-secretase inhibition by CompE increased β-CTFs expression levels by 600% in HEK APP^WT^ (Fig. 2B) while sAPPβ, Aβ_1-40_ and Aβ_1-42_ secretion was reduced by 55%, 85% and 70%, respectively (Fig. 2C-D). These results show that γ-secretase inhibition reduced β-CTFs cleavage into Aβ peptides but surprisingly also reduced sAPPβ secretion, suggesting that γ-secretase inhibition led to a reduction of β-secretase cleavage of APP.

**Figure 2:**
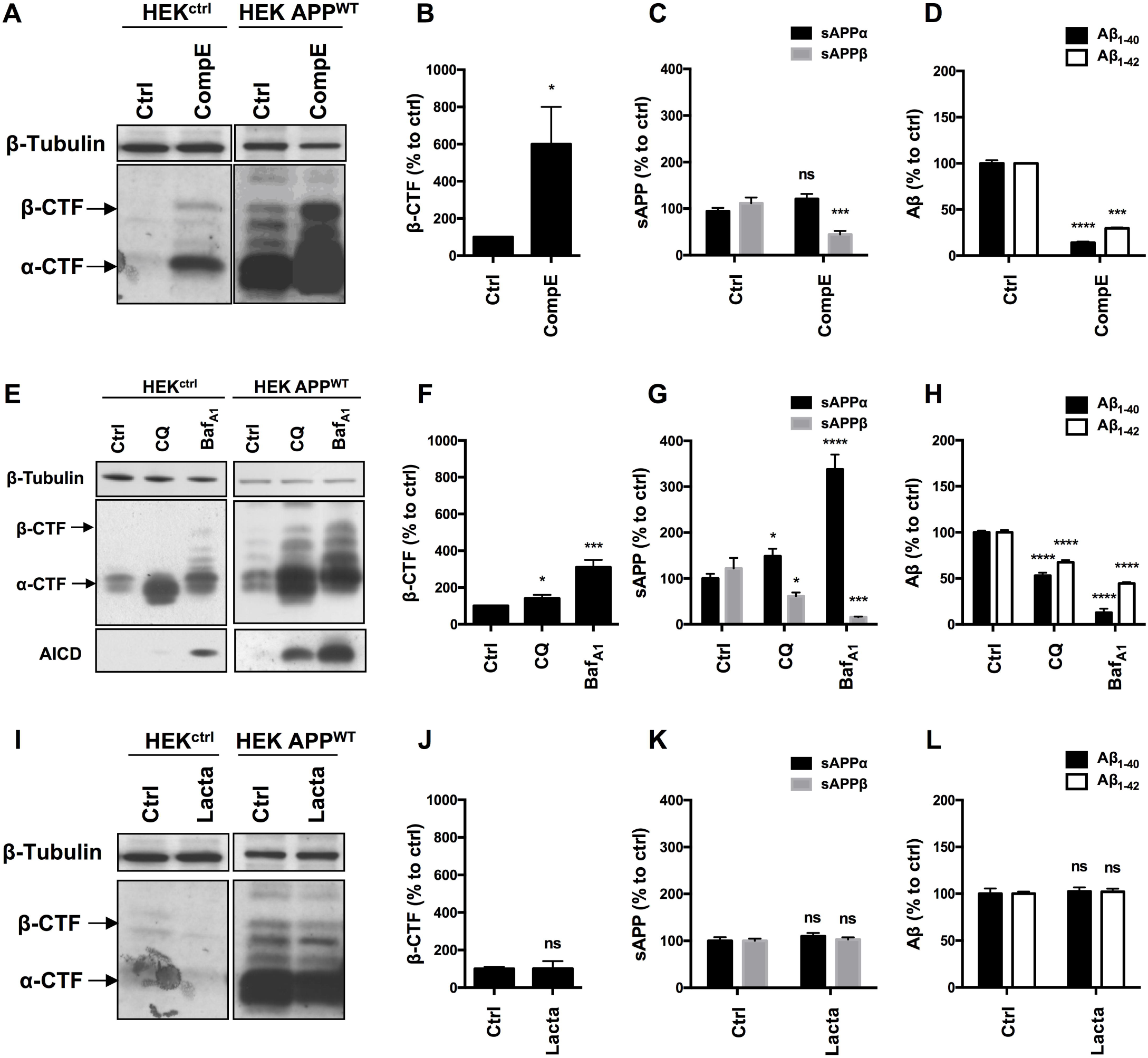
Effect of γ-secretase, lysosomal proteases or proteasome inhibition on APP processing in HEK^ctrl^ and HEK APP^WT^ cells. HEK 293 cells overexpressing or not APP^WT^were treated with Compound E (CompE, 30nM, 6h), Chloroquine (CQ, 10μ,M, 24h), Bafilomycin A1 (Baf_A1_, 100nM, 24h) or Lactacystin (Lacta, 5μM, 6h). **(A, E, I)** Western-blot analysis of β-tubulin and APP-CTFs. β-tubulin staining was used as loading control. α-CTFs and β-CTFs are indicated by arrows. **(B, F, J)** Western-blot quantification of β-CTFs from HEK APP^WT^ cells, expressed as a percentage of the control condition. **(C, G, K)** Quantification of secreted sAPPα (black bars) and sAPPβ (grey bars) by electro-chemiluminescence immunoassay from HEK APP^WT^ cells, expressed as a percentage of the control condition. **(D, H, L)** Quantification of secreted Aβ_1-40_ (black bars) and Aβ_1-42_ (white bars) measured by ELISA from HEK APP^WT^ cells, expressed as a percentage of the control condition. Data are expressed as the mean ± SEM (n=3), ^∗^p<0.05, ^∗∗^p<0.001, ^∗∗∗^p<0.001 and ^∗∗∗∗^p<0,0001.

Next, lysosomal proteases were inhibited using the alkalizing agent Chloroquine (CQ), a weak base, or Bafilomycin A1 (Baf_A1_), a vacuolar proton pump inhibitor. CQ and Baf_A1_ treatments both induced a significant increase in AICD and APP-CTFs, especially α-CTFs in both HEK^ctrl^ and HEK APP^WT^ (Fig. 2E). In HEK APP^WT^ treated cells, β-CTFs expression was increased by 40% with CQ and by 300% with Baf_A1_ (Fig. 2F). Secretion of sAPPβ was reduced by 40% with CQ and by 85% with Baf_A1_ while sAPPα was significantly increased by 49% and 230%, respectively (Fig. 2G). CQand Baf_A1_ treatments also significantly repressed Aβ_1-40_ and Aβ _1-42_ secretion (Fig. 2H). Treatments with CQ or Baf _A1_ inhibited β-CTFs degradation and reduced Aβ peptides and sAPPβ production while they increased sAPPα secretion, suggesting that the amyloidogenic pathway was reduced in favour of the α-secretase cleavage. These results are in agreement with previous reports showing that the endosomal/lysosomal pathway is of particular importance for the whole APP processing, and in particular for efficient amyloidogenic processing [14,15,17].

Finally, the proteasome involvement was assessed using the specific inhibitor lactacystin (Lacta) (Fig. 2I). Importantly, APP-CTFs levels were not significantly modified after 6h of lactacystin treatment in HEK^ctrl^ or in HEK APP^WT^ treated cells (Fig. 2I and J). Moreover, proteasome inhibition did not change the secretion of sAPPα/sAPPβ or AP_1-4_ and AP_1-42_ peptides suggesting that APP processing is not modulated by the proteasome as far as our cell system is concerned (Fig. 2K and L).

Together, these results demonstrate that γ-secretase and lysosomal pathway are directly involved in the processing of sAPP and APP-CTFs that derive from full-length APP, while the proteasome is not involved in the degradation of these fragments in our models. Importantly, similar results were obtained both in naïve HEK, expressing endogenous levels of APP, and in HEK over-expressing APP^WT^, thus showing that overexpressed APP^WT^ follows the same processing as the endogenous APP, making it a valid model to study APP processing.

### Overexpressed C99 is processed by γ-secretase and the proteasome

Given the inconsistency between our results and the literature with regards to the involvement of the proteasome towards APP processing, we used another model, which consisted on HEK cells stably expressing the last 99 carboxy-terminal residues of APP (HEK^C99^) fused to a signal peptide [30]. Indeed, most of the studies aimed at analysing the role of the proteasome in APP processing were performed in C99 overexpressing cells. In HEK^C99^, γ-secretase inhibition using CompE doubled the amount of C99 levels (Fig. 3A and 3B) and reduced by 30% the secretion of Aβ_1-40_ peptides (Fig. 3C). Aβ_1-42_ peptides could not be detected in these conditions, probably due to their low levels of production as described in earlier reports [25]. Surprisingly, in sharp contrast with results obtained in HEK APP^WT^ cells, treatments with CQ or Baf_A1_ did not significantly modify the amount of C99 (Fig. 3D-E), suggesting that overexpressed C99 is not degraded by lysosomal proteases. However, while CQ had no effect on Aβ_1-40_ peptides *secretion*, Baf_A1_ induced a 50% increase of Aβ_1-40_ peptides secretion (Fig. 3F). Finally, proteasome inhibition with lactacystin induced a significant increase of C99 compared to the control condition (Fig. 3G-H). Lactacystin also induced a 50% increase in Aβ_1-40_ peptides secretion (Fig. 3I).

**Figure 3:**
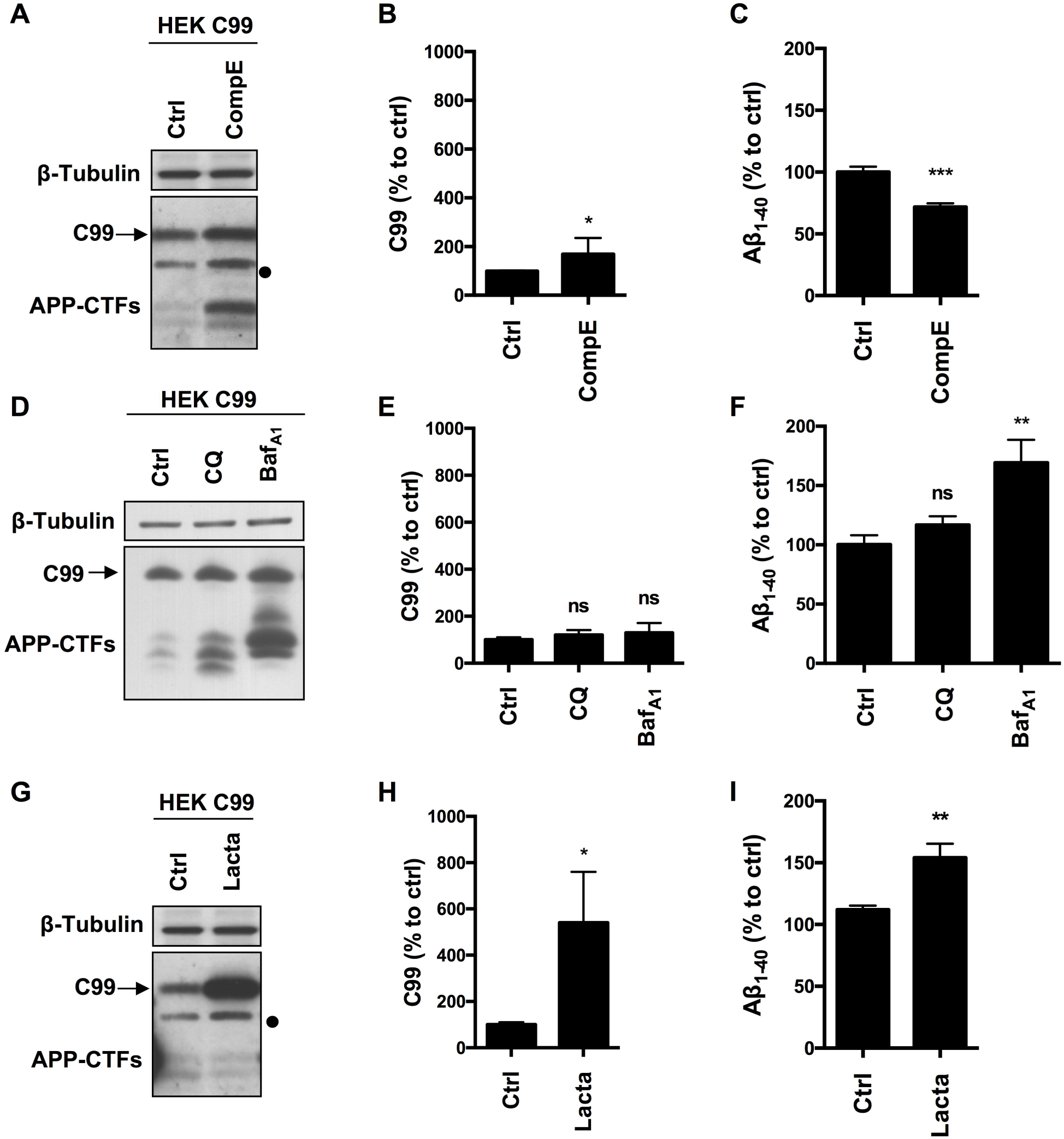
Effect of γ-secretase, lysosomal proteases or proteasome inhibition on overexpressed C99 processing in HEK cells. HEK 293 cells overexpressing C99 (SPA4CT) were treated with Compound E (CompE, 30nM, 6h), Chloroquine (CQ, 10μM, 24h), Bafilomycin A1 (Baf_A1_, 100nM, 24h) or Lactacystin (Lacta, 5μM, 6h). **(A, D, G)** Western-blot analysis of β-tubulin and APP-CTFs. **(B, E, H)** Western-blot quantification of C99 from HEK C99, expressed as a percentage of the control condition. **(C, F, I)** ELISA quantification of secreted Aβ_1-40_, expressed as a percentage of the control condition. Data are expressed as the mean ± SEM, (n=5), ^∗^p<0.05, ^∗∗^p<0.001, ^∗∗∗^p<0.001 and ^∗∗∗^∗p<0,0001. The dot on the right of Western-blot A and G is an artefactual band often revealed by the secondary antibody.

Taken together, these results show that overexpressed C99 is mainly processed by the proteasome and to a lesser extent by γ-secretase. However, in contrast with APP-derived β-CTFs, the lysosomal pathway is not involved in the degradation of overexpressed C99. Overall, these data strongly suggest that overexpressed C99 does not follow the same degradation route as the endogenously produced APβ-CTF fragments.

### Similar degradation processes are conserved in primary rat neuronal cells Infected with APP^WT^ or C99

Finally, to exclude any cell line dependent effect, we used primary neuronal rat cells that were infected or not with APP^WT^ or C99 to validate the proteasome involvement in β-CTFs processing in a more physiological model. Lactacystin treatment did not modify APP-CTFs expression in control or APP^WT^ infected neurons (Fig. 4A-B), as it was observed in HEK cells. However, overexpressed C99 was significantly accumulated in C99 infected neurons under lactacystin treatment (Fig. 4C-D). Altogether, these results showed that the proteasome is not involved in the degradation of APP-derived CTFs in cultured rat neurons, whereas proteasomal inhibition strongly repressed overexpressed C99 degradation. These data are in agreement with results obtained in HEK APP^WT^and HEK C99 cells respectively.

**Figure 4:**
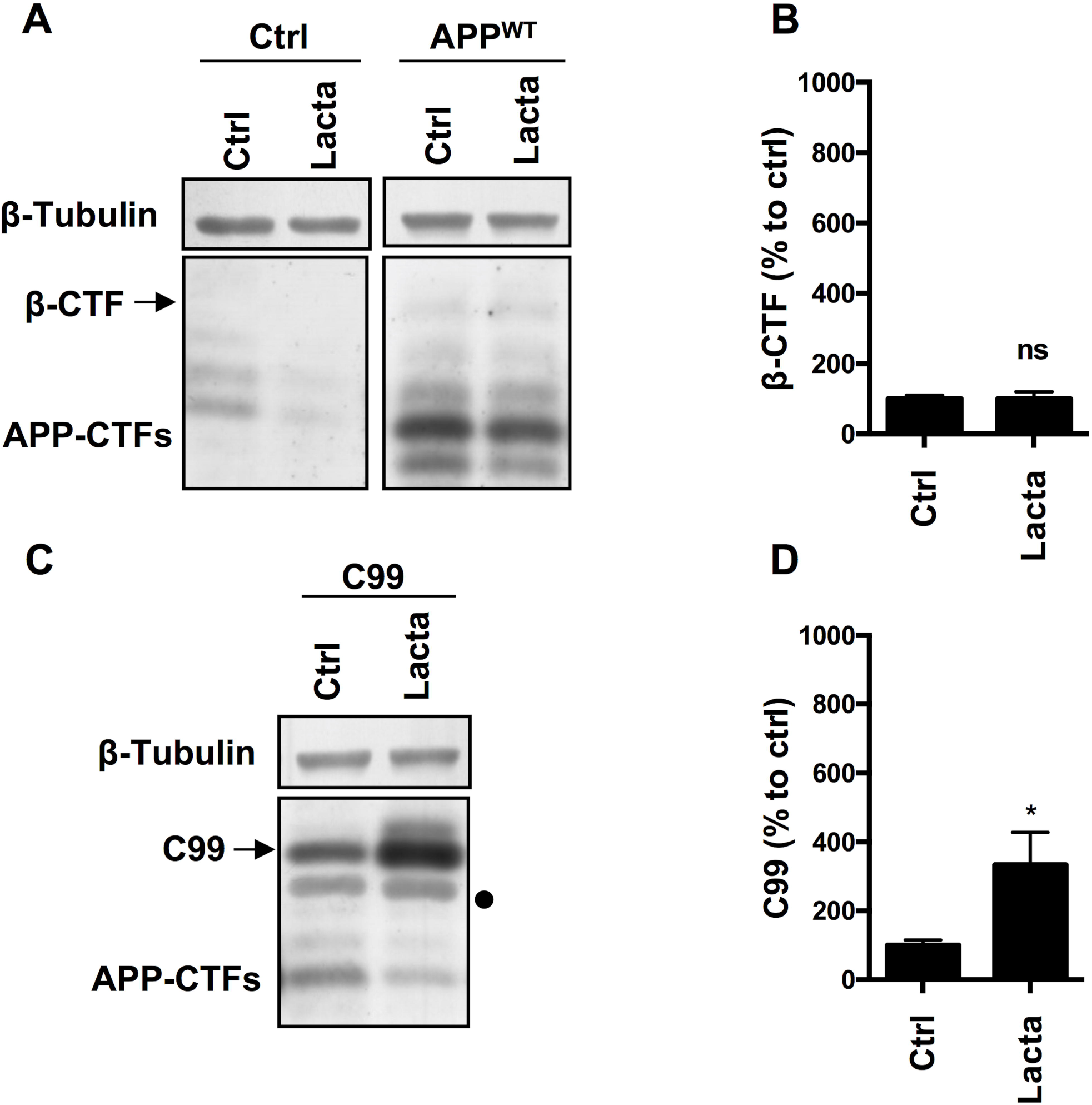
Effect of proteasome inhibition on APP-derlved CTFs and C99 in rat primary neuronal culture cells. Rat primary neuronal cultures were treated with lactacystin (Lacta, 5μM, 6h) 4 days after viral infection with either APP^WT^ or C99. **(A, C)** Western-blot analysis of β-tubulin and APP-CTFs from APP^WT^ **(A)** or C99 **(C)** infected neurons. **(B, D)** Western-blot quantification of P-CTFs and C99 from APP^WT^ **(B)** or C99 **(D)** infected neurons, expressed as a percentage of the control condition. Data are expressed as the mean ± SEM (n=3), ^∗^ p<0.05.

## Discussion

APP metabolism is a key feature in the onset and progression of Alzheimer’s disease as it is at the origin of β-CTFs accumulation and Aβ peptides production. Drugs that improve the clearance of APP metabolites would possibly protect from the detrimental effects of APP-CTFs and Aβ peptides accumulation. Therefore, a better understanding of alternative degradation pathways and molecular factors modulating β-CTFs accumulation is of interest and could uncover potential interesting pharmacological targets.

Herein, we showed that APP-CTFs produced from endogenous or overexpressed full-length APP are mainly processed by γ-secretase and the endosomal/lysosomal pathway (observations are summarized in Fig.5A). Interestingly, γ-secretase inhibition did not only lead to a reduction of Aβ peptides secretion but also to a decreased secretion of sAPPβ. This is, to the best of our knowledge, the first report suggesting that γ-secretase inhibition also repress to some extend β-secretase cleavage of APP. By which mechanism could this cross-inhibition occur? Do γ-secretase substrates include proteins that could be important for the targeting of BACE or the acidification of intracellular compartments? Is it possible that inhibiting γ-secretase activity reroute APP towards an alternative β-secretase activity, i.e. meprin β, thus producing sAPPβ that would not be recognized by the electrochemiluminescence immunoassay kit used in this study? These intriguing questions remain open and require further investigations. Our results also confirmed that besides being cleaved by γ-secretase, APP-CTFs are efficiently degraded by the endosomal/lysosomal pathway, as it was previously reported [14,15,17]. Along with reducing APP-CTFs degradation, drugs that were used to inhibit the endosomal/lysosomal pathway significantly reduced the secretion of Aβ peptides and sAPPβ species. As we previously showed that the inhibition of the endosomal/lysosomal pathway does not alter γ-secretase activity [16], these results suggest that there could be an impairment of the extracellular release of Aβ and sAPPβ from recycling compartments or that the β-secretase cleavage of APP is inhibited by these lysomotropic agents. The cleavage of APP by BACE1 occurs in acidic endosomal compartments, with an optimal activity at pH 4.5 [31,32]. Thus, it is not surprising that these drugs, which act by reducing the acidification of the endosomal and lysosomal compartments, would lead to an inhibition of BACE1-mediated APP processing and a parallel increase of sAPPα secretion, which could result from a redirection of APP to the non-amyloidogenic pathway. The development of drugs/molecules that derive from the CQ. structure and share its effects on APP processing is an interesting strategy that is currently developed to tackle AD [33].

**Figure 5:**
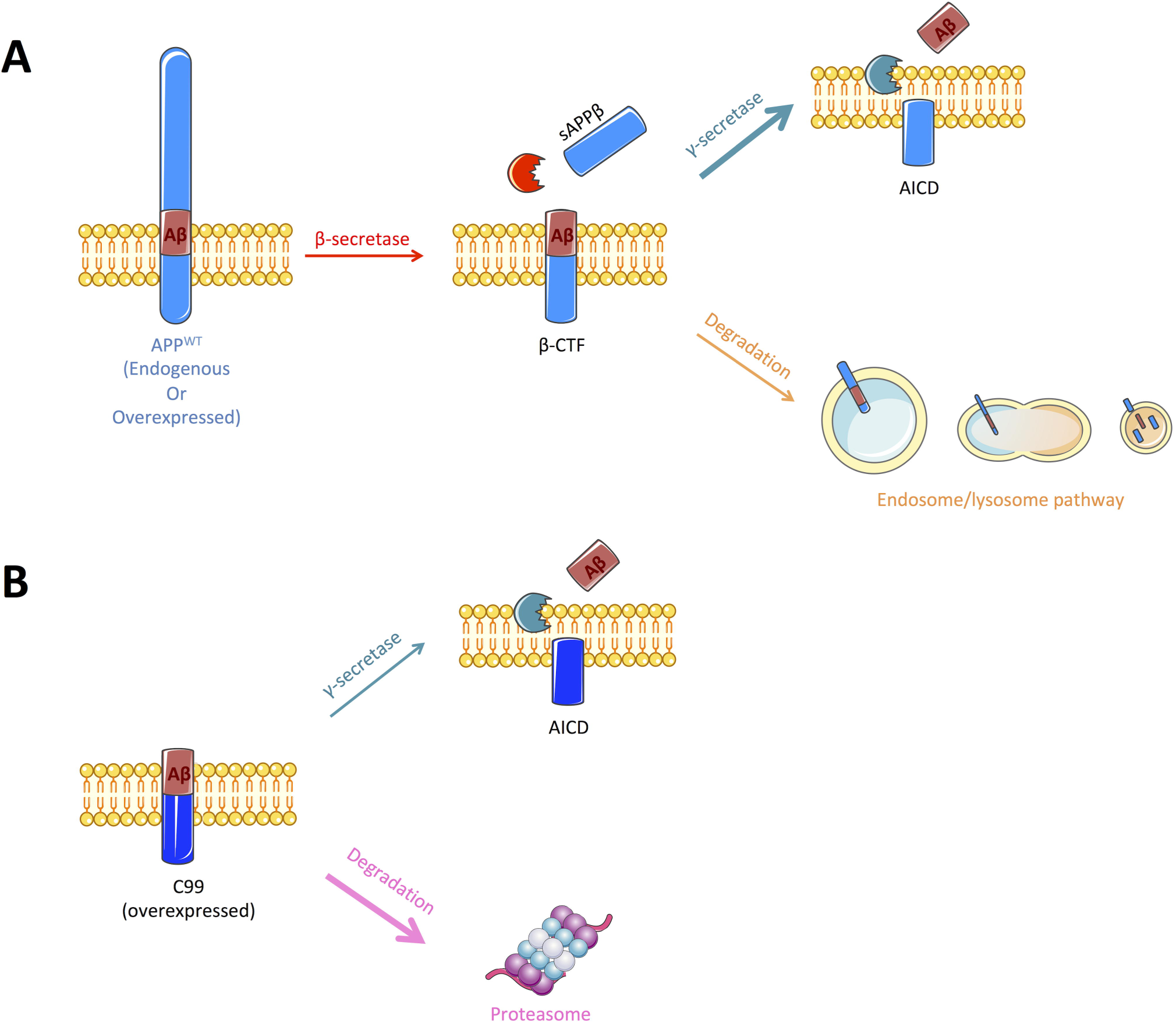
Schematic representation of the pathways implicated In the cleavage or degradation of endogenous APP-derived β-CTF or overexpressed C99. **(A)** β-CTF produced from β-secretase cleavage of full-length APP is mainly cleaved by the γ-secretase to produce Aβ peptides (nearly 60%). Remaining β-CTFs (40%) are degraded by the endosomal/lysosomal pathway. **(B)** Overexpressed C99 can also be degraded by the γ-secretase to produce Aβ peptides, but only 30% are processed through this pathway. The remaining 70% are degraded by an alternate proteasome-dependant pathway. Figure was produced in part using Servier Medical Art.

In sharp contrast to APP derived β-CTFs degradation, we found that overexpressed C99 was mainly degraded by the proteasome and to a lower extent by γ-secretase. Indeed, lactacystin led to a strong accumulation of overexpressed C99 that was accompanied by an increase in Aβ_1-40_ peptides secretion. The strong accumulation of C99 following proteasome inhibition could lead to an upsurge in γ-secretase substrate availability and therefore to an increase in Aβ peptides production, without affecting γ-secretase activity. However, we cannot exclude the possibility that γ-secretase activity could be altered following proteasome inhibition. Indeed, components of the γ-secretase complex have been shown to be ubiquitinylated and thus suggested to be handled by the proteasome [34–37].

The different processing of overexpressed C99 as compared to APP-derived β-CTFs by the proteasome could be an artefactual consequence of the overexpression. Indeed, several reports have shown that overexpressed proteins can be processed in an artefactual way by the proteasome [38]. As full-length APP was also overexpressed in our experiments, we verified whether proteasome inhibitors modified or not full-length APP expression. Our results demonstrated that full-length overexpressed APP is not sensitive to pharmacological inhibition of the proteasome in any of the cellular models analysed (data not shown). The most straightforward conclusion here is that, when ectopically over-expressed, C99 is recognized as a truncated membrane protein that will undergo ER-associated degradation (ERAD) mediated by the proteasome [39]. This is not the case for APP CTFs that are endogenously produced (and not overexpressed) along the late endosome/lysosome pathway. This is supported here by the fact that overexpressed C99 is not alternatively degraded by lysosomes, indicating that it has different subcellular location than APP-derived CTFs.

In conclusion, our data show that APP-derived CTFs are processed principally by γ-secretase and alternatively degraded by lysosomes. Indeed, the physiological processing of APP by α- or β-secretase relies on the endosomal/lysosomal pathway, which dysfunction leads to a complete disturbance of APP metabolites. However, a direct degradation of APP-derived CTFs by a proteasome-dependent pathway is not supported experimentally in this study. In sharp contrast, C99 chimeric construct is mainly processed by a proteasome-dependent mechanism and to a lesser extent by γ-secretase. Hence, homologies between APP-derived β-CTFs and overexpressed C99 processing are likely restricted to their processing by γ-secretase, therefore caution should be taken when using this model to study β-CTFs biology.

## Acknowledgments

We thank Pr. F. Checler for kindly providing us with cell lines. This work was supported by the ANR VIDALZ (ANR-15-CE18-0002) and the Labex DISTALZ (Development of Innovative Strategies for a Transdisciplinary Approach to Alzheimer’s disease). Caroline Evrard holds a doctoral scholarship from Lille 2 University.

## Conflict of interest

The authors declare that they have no conflict of interest.

